# Detection of functional activity in brain white matter using fiber architecture informed synchrony mapping

**DOI:** 10.1101/2022.02.23.481698

**Authors:** Yu Zhao, Yurui Gao, Zhongliang Zu, Muwei Li, Kurt G. Schilling, Adam W. Anderson, Zhaohua Ding, John C. Gore

**Author notes:** The corresponding authors: Zhaohua Ding and Yu Zhao ( and).

## Abstract

A general linear model is widely used for analyzing fMRI data, in which the blood oxygenation-level dependent (BOLD) signals in gray matter (GM) evoked in response to neural stimulation are modeled by convolving the time course of the expected neural activity with a canonical hemodynamic response function (HRF) obtained a priori. The maps of brain activity produced reflect the magnitude of local BOLD responses. However, detecting BOLD signals in white matter (WM) is more challenging as the BOLD signals are weaker and the HRF is different, and may vary more across the brain. Here we propose a model-free approach to detect changes in BOLD signals in WM by measuring task-evoked increases of BOLD signal synchrony in WM fibers. The proposed approach relies on a simple assumption that, in response to a functional task, BOLD signals in relevant fibers are modulated by stimulus-evoked neural activity and thereby show greater synchrony than when measured in a resting state, even if their magnitudes do not change significantly. This approach is implemented in two technical stages. First, for each voxel a fiber-architecture-informed spatial window is created with orientation distribution functions constructed from diffusion imaging data. This provides the basis for defining neighborhoods in WM that share similar local fiber architectures. Second, a modified principal component analysis (PCA) is used to estimate the synchrony of BOLD signals in each spatial window. The proposed approach is validated using a 3T fMRI dataset from the Human Connectome Project (HCP) at a group level. The results demonstrate that neural activity can be reliably detected as increases in fMRI signal synchrony within WM fibers that are engaged in a task with high sensitivities and reproducibility.

## I. Introduction

**F**UCTIONAL MRI (fMRI) based on blood oxygenation level-dependent (BOLD) signals is well established for detecting and mapping neural activities in brain gray matter (GM). However, there are relatively few corresponding reports of fMRI studies of white matter (WM), partly due to continuing uncertainties regarding the source of BOLD signals therein ^1,2^ as well as technical challenges in their reliable detection. Nonetheless, the existence and potential functional relevance of BOLD signals in WM have been validated by a growing number of recent studies, (see reviews in ^3,4^). These have been partially motivated by the fact that WM comprises about half of the brain parenchyma and plays an important role as the communication system of the brain, and so the functional properties of WM should be included in any complete model of brain organization. In the emerging field of WM fMRI, analysis methods established for GM have been adopted and applied, including measures of function connectivity ^5-7^ and spectral analysis ^8-11^, and these have shown the potential of characterizing selected features of WM in neurodegenerative and other disorders.

The standard approach that has been widely used in fMRI studies of GM uses a general linear model (GLM) to detect activations that correspond to a change in the local BOLD signal magnitude, but this may not be appropriate for WM as suggested by previous reports ^12^. The GLM assumes that the BOLD response to stimulation may be treated as a linear shift-invariant system so that the BOLD signals can be modeled by convolving the time course of neural activities with a hemodynamic response function (HRF). The GLM assumes that evoked neural activations are synchronous with the timings of presented stimuli or tasks that trigger hemodynamic responses ^13^ and correspond to a change in signal magnitude. This assumption may not be valid in all situations, especially in WM because it ignores intrinsic brain activities that pre-exist in the background and are unrelated to extrinsic stimuli. Such intrinsic brain activities were considered noise before they were found to predict spatial patterns of functional connectivity (i.e., functional networks) in a resting state ^14,15^. A voxel in WM contains a large number of axons that connect neurons in different positions in GM ^16,17^ and these may produce asynchronous neuronal signals so that fMRI signals measured in a WM voxel may be more complex than those in GM and thus inconsistent with stimuli in tasks. A further assumption in the GLM is that the HRF keeps fixed for all positions, as implemented in SPM ^13^. This assumption is not valid in all WM as previous studies have shown that HRFs measured in WM exhibit substantial regional variations ^12,18^. A potential solution to this issue is to use variants of a GLM that allow HRFs to adapt spatially. However, potential mismodeling of HRF in such approaches may prevent accurate activation mapping and reduce sensitivity. Furthermore, fMRI signal fluctuations in WM are substantially lower than those in GM ^19^ due to the significantly lower blood volume in WM compared to GM ^1,20^ and the resulting lower BOLD signal-noise-ratio poses additional challenges to the analysis of WM.

A model-free approach, without the assumptions mentioned above, that produces robust detection of activity-related BOLD signals in WM is thus needed. An alternative to the GLM that incorporates the temporal coherence of a voxel with respect to its neighbors may provide a means of activation mapping without modeling the BOLD signal or assuming local changes in signal magnitude. This approach is inspired by previous findings that the regional homogeneity (ReHo) of fMRI signals in GM is modulated by stimuli presented in tasks ^21-23^. The ReHo approach relies on the fact that voxels within a GM volume at a macroscopic scale constitute a functional unit in which the synchrony of inter-voxel time courses can be enhanced under functional loading. In contrast to GM, anatomical structures of the WM feature complex interwoven networks, in which fibrous tissues constitute elongated functional units for neural signal transmission and thus show structural anisotropy. Evidently, brain functional processes embedded in such an anisotropic substrate should also exhibit anisotropic spatial profiles. This has been demonstrated by previous studies of BOLD signals in WM ^24-26^, which reported structure-specific temporal correlations along WM tracts in a resting state that become more pronounced in relevant structures under a task loading. Given the relationships between local fiber architectures and WM function, we propose a new approach for mapping activity-related BOLD signals in WM that incorporates a metric of BOLD signal synchrony. This approach introduces two technical innovations. First, an adaptive spatial window is designed for each voxel to extract and weight time courses associated with a fiber-architecture-informed neighborhood of the voxel. The neighborhood is determined using diffusion orientation distribution functions (ODFs) obtained from high angular resolution diffusion imaging (HARDI) data. A similar concept was proposed in a previous report that used anisotropic spatial filtering to enhance noisy BOLD contrasts in WM ^27^. Second, a modified principal component analysis (PCA) is applied to time courses of fMRI data to estimate the synchrony of time courses within the fiber-architecture-informed window of each voxel. Taken together, the proposed approach overcomes the intrinsic limitations of GLM based analyses of BOLD signals in WM.

In the following, the proposed model-free mapping of neural activations is illustrated using a 3T fMRI dataset from the Human Connectome Project (HCP) at a group level. First, spatial patterns within activation maps are evaluated by reference to anatomical structures of WM fibers derived from diffusion MRI. Then, the influence of task loading on activation are investigated by comparing activation maps derived from motor-task fMRI data with those acquired in a resting-state. Motor-task enhanced activations in WM are demonstrated in WM regions related to the primary motor cortex, or M1, which are determined by WM fiber tracking with seed points in M1. Furthermore, the sensitivity, specificity and reproducibility of the proposed method are assessed statistically based on fiber-tracking-based parcellations of task-related and unrelated WM regions.

## II. Methods

The proposed approach consists of two major steps implemented on a voxel-by-voxel basis (**Fig. 1**). First, a fiber-architecture-informed spatial window is created for each voxel using ODFs constructed from HARDI data. Second, a modified PCA is used to estimate the synchrony of fMRI time courses based on this spatial window. Detailed algorithms and procedures of the proposed approach are described as follows.

**Fig. 1.**
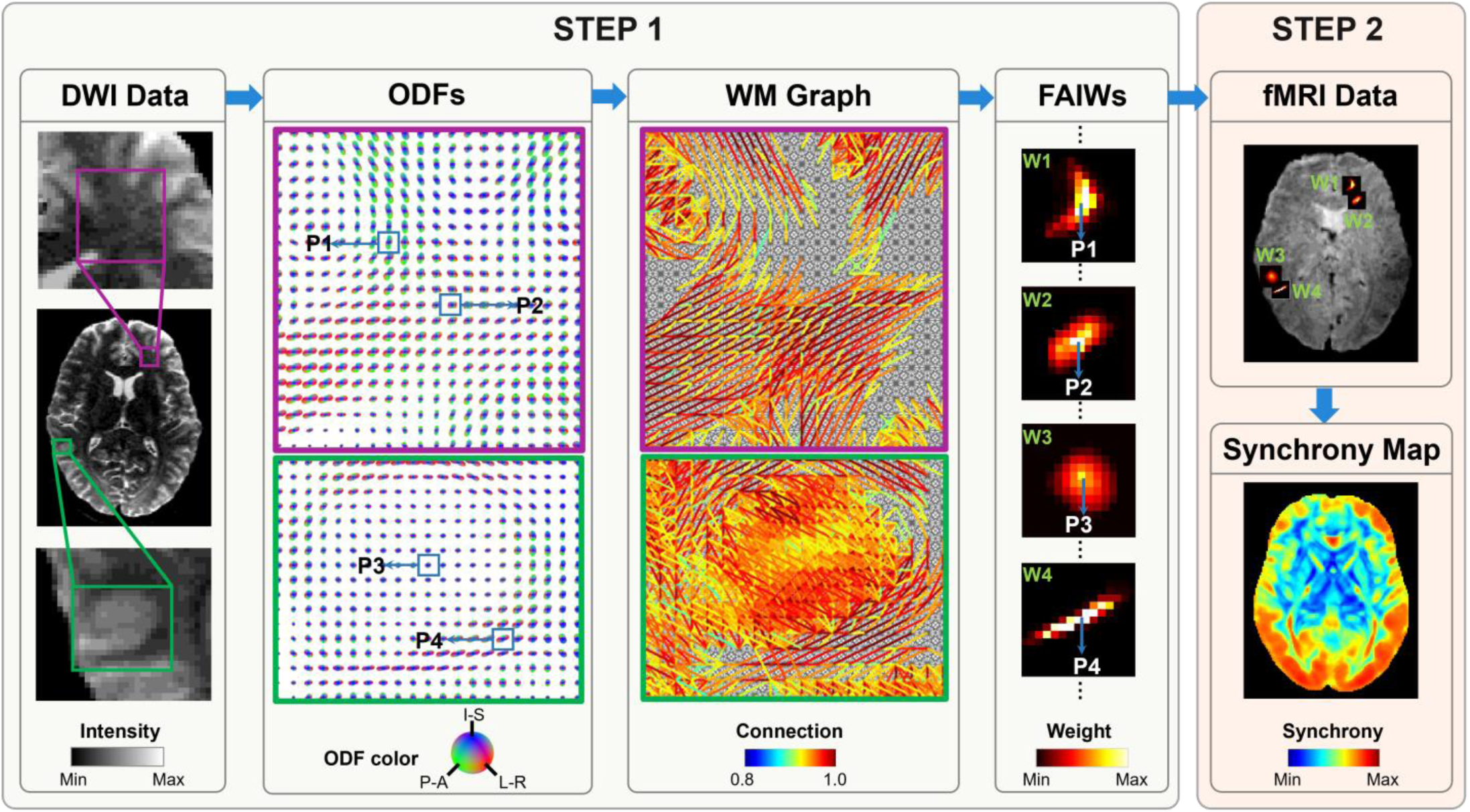
Schematic of fiber architecture informed synchrony mapping in human white matter (WM). The algorithm of the proposed approach consists of two major steps. First, a spatial window is created for each voxel based on local fiber architectures. In this step, orientation distribution functions (ODFs) constructed from diffusion MRI data provide the information on fiber architectures, which are then used to generate a topological graph for WM. In this graph, every vertex corresponds to a voxel in MR images, with weighted connections to its neighbors. The edge weights are determined by the coherence between the directions of diffusion and the orientation of the graph edges. Then, diffusion on the graph with a point source located at each vertex is simulated to produce a diffusion profile that can be further used as a fiber-architecture-informed window (FAIWs). Here, typical FAIWs created for four voxels (**P1, P2, P3** and **P4**) demonstrate that they are adaptive to local fiber architectures. Second, a modified PCA is implemented to estimate the synchrony of fMRI time courses based on the FAIWs from the first step to yield activation maps. In the modified PCA, the FAIWs emphasize the time courses associated to voxels near the central position with high weights, which ensures the spatial specificity of the synchrony estimate.

### A. Design of fiber-architecture-informed spatial window

Principles from the recently emerging field of graph signal processing (GSP) ^28,29^ provide an elegant framework to design the fiber-architecture-informed spatial window. Based on the GSP framework, we analyze fMRI time courses at a discrete set of positions, which form a set of vertices of a topological graph, in such a way that underlying constraints from fiber architectures can be accounted for by edges of the graph. Here, we denote a graph as **G** = {**V, E, W**} as usual, which consists of a finite set of vertices **V** with |**V**| = **N**, a set of edges **E**, and a weighted connection matrix **W. W**_***i***,*j*_ represents the connection strength of edge ***e*** = (*i, j*) that connects vertices *i* and *j*. On the vertices of the graph, we define a vector **f** ∈ ℝ^*N*^ to represent a set of auxiliary quantities to be used to generate spatial windows, where the *i*-th component of the vector, **f**(***i***), represents the value at the *i*-th vertex in **V**.

For this study, voxel-wise graphs are constructed using ODFs derived from HARDI data, which have been well described in previous reports ^27,30^. In each graph, every vertex corresponds to a voxel in 3D MR images with weighted connections to its neighbors. The connection weight of any two neighboring voxels depends on the coherence of diffusion orientations between the two voxels. As a result, two neighboring voxels residing in the same fiber tract have relative larger weight than those in different fiber tracts.

Building on the topological properties of the fiber-architecture-informed graph, a spatial window is created for each vertex/voxel by simulating diffusion on the graph with point sources located at each vertex. A kernel is defined for the diffusion process in the spectral domain, which is derived from a Laplacian matrix defined in GSP. Specifically, a Laplacian matrix of **G** is defined as **L** = **D** − **W**, where **D** denotes the graph’s degree matrix, which is diagonal with elements ***d***_***i***,***i***_ = Σ_***j***_ **w**_***i***,***j***_; the graph operator **L** is a real symmetric matrix, and thus has a complete set of orthonormal eigenvectors {***u***_*𝓁*_} _*𝓁*=1,2,…,*N*_ and associated eigenvalues {***λ***_*𝓁*_} _𝓁=1,2,…,*N*_ satisfying **L*u***_𝓁_ = *λ*_𝓁_***u***_𝓁_. With these eigenvectors, the graph signal **f** can be transformed into the spectral domain,

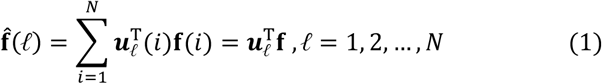

This spectral representation allows an exact reconstruction, i.e., the signal **f** can be recovered as

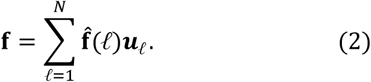

In the graph spectral domain, the kernel of diffusion ^31^ can be defined as

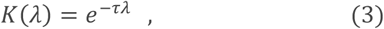

where *τ* is a free parameter determining the spatial extent of diffusion on a graph, and it can be intuitively treated as the duration of diffusion. Given an initial point source **f**_0_ (t=0), the transient diffusion profile at time point t can be derived as

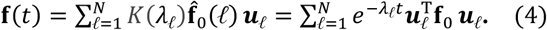

To create a spatial window centered at a voxel *i* we define this point source by setting the initial profile **f**_0_ to be equal to 1 at the position ***i*** and 0 otherwise. The transient profile **f**(*t*) at the time point t is further processed to yield the fiber-architecture-informed spatial window, the width of which can be adjusted by the parameter *τ*. To be specific, the window is defined by taking voxels with *M* largest values of **f**(*t*), which is subsequently normalized with their summation,

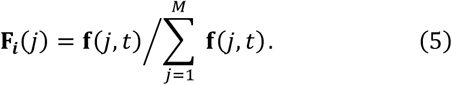

In this study, the largest parameter *M* is taken under an empirical constraint 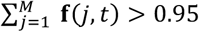, wherein small values in the diffusion profile of each voxel are removed to improve computational efficiency.

### B. PCA based on fiber-architecture-informed window

In previous reports, the synchrony of fMRI time-courses in a parcellated region have been quantified using PCA ^32^, wherein the variance explained by the first principal component gauges the extent of synchrony. Here, the conventional PCA is modified to allow the integration of fiber architecture information, referred to as a modified PCA based on a fiber-architecture-informed window (FAIW-PCA). Basically, the raw fMRI time courses are first normalized with respect to the mean values and the standard deviations. Then, within each window F_***i***_ centered at the position i, the normalized time courses are rearranged into a matrix (X) of dimension T × M, where T denotes the number of time frames and M the number of voxels within the window. We denote a time course in X as ***x***_***i***_, the i-th column in the matrix X, and formulate the FAIW-PCA as an optimization problem, where the first principal component *v*_1_ can be obtained by

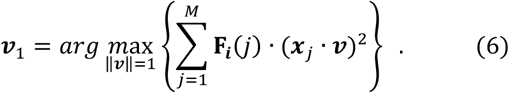

Note that, without the weights **F**_***i***_(*j*), this optimization problem will degenerate into a conventional PCA. Here, **F**_***i***_(*j*) is applied in the optimization such that the time courses associated with the voxels near the center position are emphasized more in the first principal component obtained, which emphasizes the spatial specificity of the synchrony estimate. We further write this optimization in matrix form,

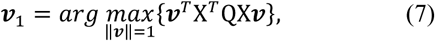

where the matrix Q = diag[(**F**_***i***_(1), **F**_***i***_(2), …, **F**_***i***_(*M*)]. With simple algebraic manipulations, Equation 7 can be written as

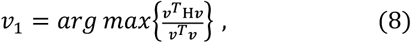

where H = X^*T*^QX. H is a positive-definite matrix, and thus has a complete set of orthonormal eigenvectors {*ν*_𝓁_}_𝓁=1,2,…,*M*_ and associated eigenvalues {σ_𝓁_} _𝓁=1,2,…,*M*_, σ_𝓁_ ≥ σ_𝓁+1_ ≥ 0 that satisfy **H***ν*_𝓁_ = σ_𝓁_*ν*_𝓁_. The quantity to be maximized in Equation 8 is a Rayleigh quotient. The quotient’s maximum possible value is the largest eigenvalue σ_1_, which is associated with the solution ***ν***_1_. Finally, the synchrony of time courses is estimated by

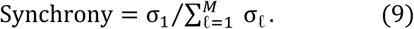

Note that, as 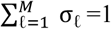 due to the normalization of the window **F**_***i***_(*j*), the synchrony can be simply estimated by σ_1_, σ_1_ ∈ [0,1].

### C. Data and preprocessing

#### Datasets

Human MRI datasets used in this study were sourced from the WU-Minn Human Connectome Project (HCP) database ^33,34^, which included a 128 subject sub-group (54% female, age range =22-37) that passed image quality control criteria and completed all imaging acquisitions. The datasets comprised resting-state fMRI, motor-task fMRI, T1-weighted MRI and diffusion-weighted MRI (DWI) and were downloaded from the HCP repository as minimally preprocessed data. The HCP data acquisition was approved by the Washington University Institutional Review Board and informed consent was obtained from all subjects. Parameters of MRI acquisitions and the minimal preprocessing are fully described in reference ^35^. The dataset was separated into two groups: the first 64 subjects for a primary dataset and other 64 subjects for an independent validation dataset.

#### Preprocessing

The HCP preprocessed functional and diffusion data are in different neurological spaces (fMRI data in ACPC space; and DWI data in MNI space). Reasonably accurate co-registration of images between the two MRI modalities, which typically involves nonlinear transformations, is critical for the proposed approach. In this work, the HCP preprocessed fMRI volumes were nonlinearly registered into ACPC space, and then up-sampled to the voxel resolution of the DWI data (i.e., 1.25mm isotropic). This spatial transformation was performed by leveraging the mni2acpc.nii displacement maps provided within the HCP preprocessed data. Note that the ACPC space was chosen as the working space in this study to avoid a spatial transformation on DWI data that would necessitate a complex adjustment of axonal orientations ^36^. In addition, the segmentation volumes (aparc+aseg.nii) provided within the HCP data were down-sampled to the working resolution, from which a WM mask was created. Further preprocessing of fMRI data in this study included four steps as follows. First, cerebrospinal fluid (CSF) nuisance signals and 12 variates of head motion were regressed out from time courses of fMRI data. Second, the volumes of fMRI data within the WM mask were spatially smoothed with a Gaussian smoothing kernel of 3-mm full width at half maximum. Spatial smoothing this way could prevent signal contamination from nearby GM. Note that, besides improving the signal-to-noise ratio in BOLD time courses, the spatial smoothing can reduce the sensitivity of the proposed approach to potential co-registration errors at some cost of spatial specificity. Third, time courses were bandpass filtered to retain frequencies from 0.01 to 0.12 Hz. Finally, the means of the filtered time courses were subtracted, and the data then normalized to unit variance.

### D. Synchrony mapping using FAIW-PCA

Synchrony mapping using FAIW-PCA was performed on the HCP image data that had been co-registered in ACPC space. To begin with, a whole brain graph was constructed for each subject using the afore-mentioned approach with HARDI data. Following reference ^27^, two tuning parameters {α, β} were set to {0.9, 50} in the generation of the graph so that the nature of elongated fibers can be characterized with high fidelities. With the graph constructed, the two-step activation mapping described in **Sections II(A)** and **II(B)** was performed voxel-wise: (1) a local fiber-architecture-informed window was created for each voxel by the spectral kernel approach, and (2) FAIW-PCA was performed to estimate the synchrony of fMRI time-courses on the basis of fiber architecture information within the window. The parameter of the kernel *τ*, which determines the spatial extent of signal diffusion on fibers, was optimized to ensure the distribution of the synchrony σ_1_ (σ_1_ ∈ [0,1]) in WM was peaked around 0.5.

### E. Validation of FAIW-PCA-based activation mapping

We validated the proposed approach at a group level in MNI space. To do so, synchrony maps computed from each subject in the primary dataset and the independent validation dataset were co-registered into MNI space by leveraging the acpc2mni.nii displacement maps provided within the HCP preprocessed data, and subsequently averaged to yield group-average synchrony maps. Then, to characterize the task-induced changes in synchrony, difference maps were calculated as

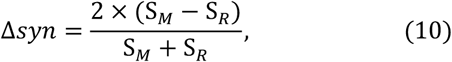

where S_*R*_ and S_*M*_ are the synchronies measured in resting state and motor task, respectively; values of Δsyn indicate the strength of neural responses to the task. The group-average synchrony maps and their difference maps estimated from the primary dataset were analyzed as follows: (1) the group-average synchrony maps from resting states were investigated first by reference to anatomical structures conveyed by T_1_-weighted (T_1_w) images and fractional anisotropy maps derived from the HARDI data. (2) fiber tracking was implemented in the DSI Studio software package ^37^ to extract WM regions related to the primary motor cortex (M1) and the auditory cortex (A1&A2); the synchrony maps from the two functional states were compared with an emphasis on the M1 related WM regions; the auditory cortex is mainly associated with processing auditory information, which is in principle not involved in motor tasks and expected to undergo minimal variations between the two functional states; thus, the specificity of the proposed approach was assessed by comparing the increase in WM activations under the motor task in the task-related and unrelated regions. (3) the regional synchronies were compared between the two functional states using the HCP tractography atlas of WM available on the website: http://brain.labsolver.org/diffusion-mri-templates/tractography. (4) group level statistical maps of Δsyn-based activation were compared with those of GLM-based activation; the statistical maps of Δsyn-based activation were derived by voxel-wise paired t-tests of differences between resting state and motor task with a false discovery rate (FDR) correction; the statistical maps of GLM-based activation were estimated using the standard SPM toolbox. Finally, comparisons of Δsyn-based activation maps between the primary dataset and the independent validation dataset were performed to validate the reproducibility of the proposed approach.

## III. Results

**Fig. 2** shows group-average synchrony maps reconstructed with the motor-task fMRI dataset collected from 64 participants in HCP, and FA color maps reconstructed with the DTI dataset from a representative participant as well as T_1_w images. Many continuous structures can be observed with high synchrony in both GM and WM. In the GM, the synchronous structures agree grossly with anatomical structures identified in the T_1_w images. In the WM, synchronous structures exist almost exclusively in regions that contain large axonal fascicles, with some typical examples denoted by red arrows that exhibit uniform fiber orientations in the FA color maps. These spatial patterns in WM are reasonable in terms of their relationship to functional neuroanatomy. For instance, the fiber bundles in occipital WM convey mainly visual signals so that voxels within them tend to have synchronous BOLD signals. Conversely, the WM regions that include fibers with heterogenous orientations (indicated by heterogeneities in FA colors) or crossing fibers (visualized as dim FA colors) exhibit low values in the synchrony maps, which can be appreciated in the regions denoted by green arrows in **Fig. 2**. This is explainable by the fact that the fibers leading to different locations in GM tend to have asynchronous neural activities and thus asynchronous BOLD signals.

**Fig. 2.**
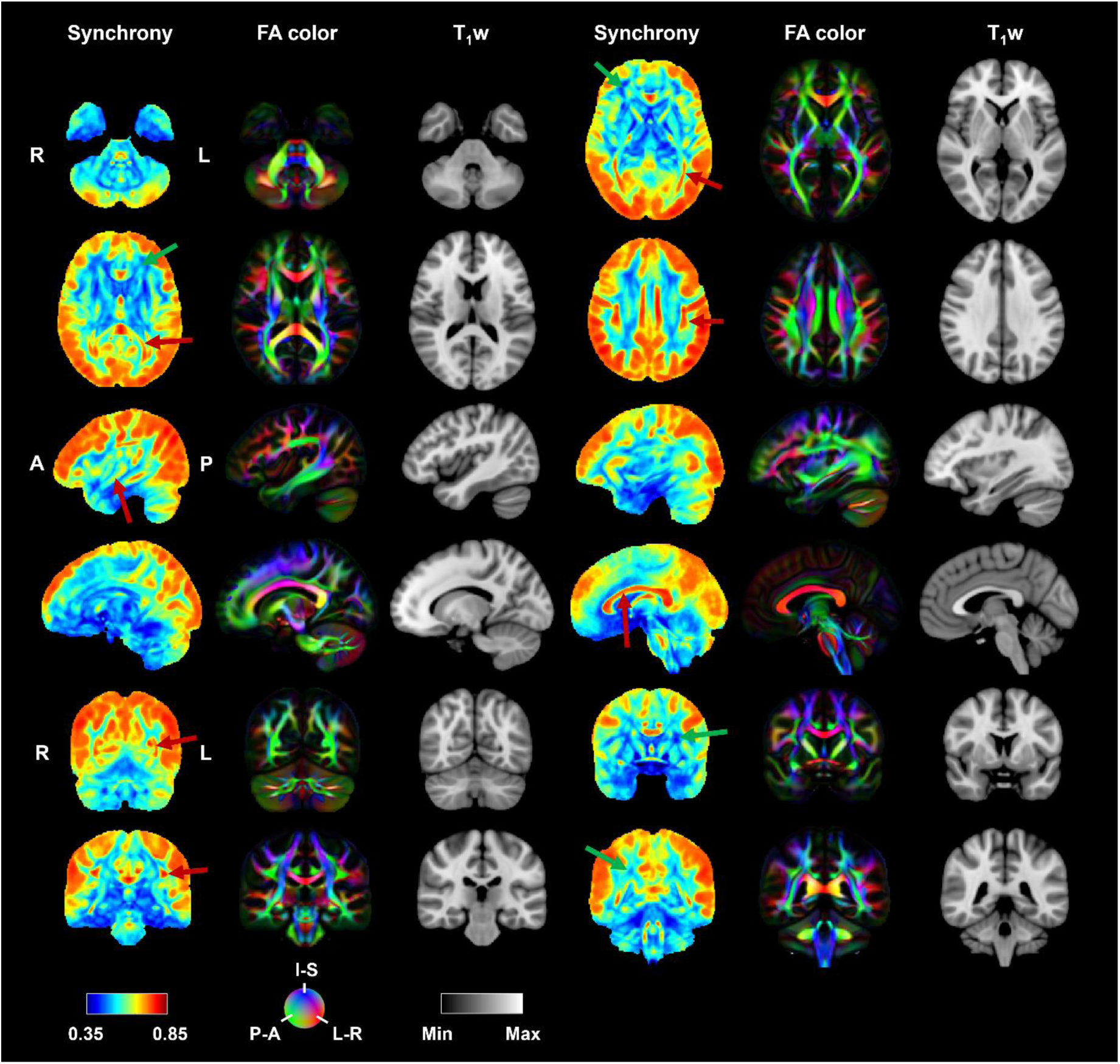
Group-average synchrony maps reconstructed with the motor-task fMRI dataset collected from 64 participants in HCP, and FA color maps reconstructed with the DTI dataset as well as T_1_w images from a representative participant. There are many structures with high synchrony in both GM and WM. In the GM, the synchrony structures agree grossly with anatomical structures identified in the T_1_w images. In the WM, synchrony structures exist almost exclusively in regions that contain large axonal fascicles, with some typical examples denoted by red arrows that exhibit uniform fiber orientations in the FA maps. Conversely, the WM regions that include fibers with heterogenous orientations (indicated by heterogeneities in FA colors) or crossing fibers (visualized as dim FA colors) exhibit low intensities in the synchrony maps, which is showcased in the regions denoted by green arrows.

**Fig. 3** compares the group-average synchrony maps estimated from the resting-state and motor-task fMRI data on the basis of WM tractograms, in which the motor-task-related brain activations can be visually appreciated. In **Fig. 3A&B**, M1 and A1&A2 related WM regions were determined by using fiber tracking with seed points defined in the corresponding cortical regions. It can be seen that, in both M1 GM regions and the related WM regions, the synchrony maps exhibit obviously higher intensities during the motor task than in the resting-state. Coactivations of the associated GM and WM regions are quantitively visualized by the maps of synchrony difference (Δsyn) between the two states in the bottom row of **Fig. 3C**. In addition, some WM regions connected with the occipital and frontal lobes also exhibit obvious enhancements of synchrony under the motor task whereas the WM regions associated with the auditory cortex have no significant changes between the two states. The changes in synchrony in M1 and A1&A2 related WM regions are further compared in **Fig. 4**, in which the synchronies during the motor task are plotted against those from the resting state. The offset distances of points in the scatter plots from the identity line reflect changes in WM voxels. Evidently, the changes in BOLD signals in the M1-related region are significantly larger than those of the A1&A2 related WM region, in which the scatter points are tightly and symmetrically distributed along the identity line.

**Fig. 3.**
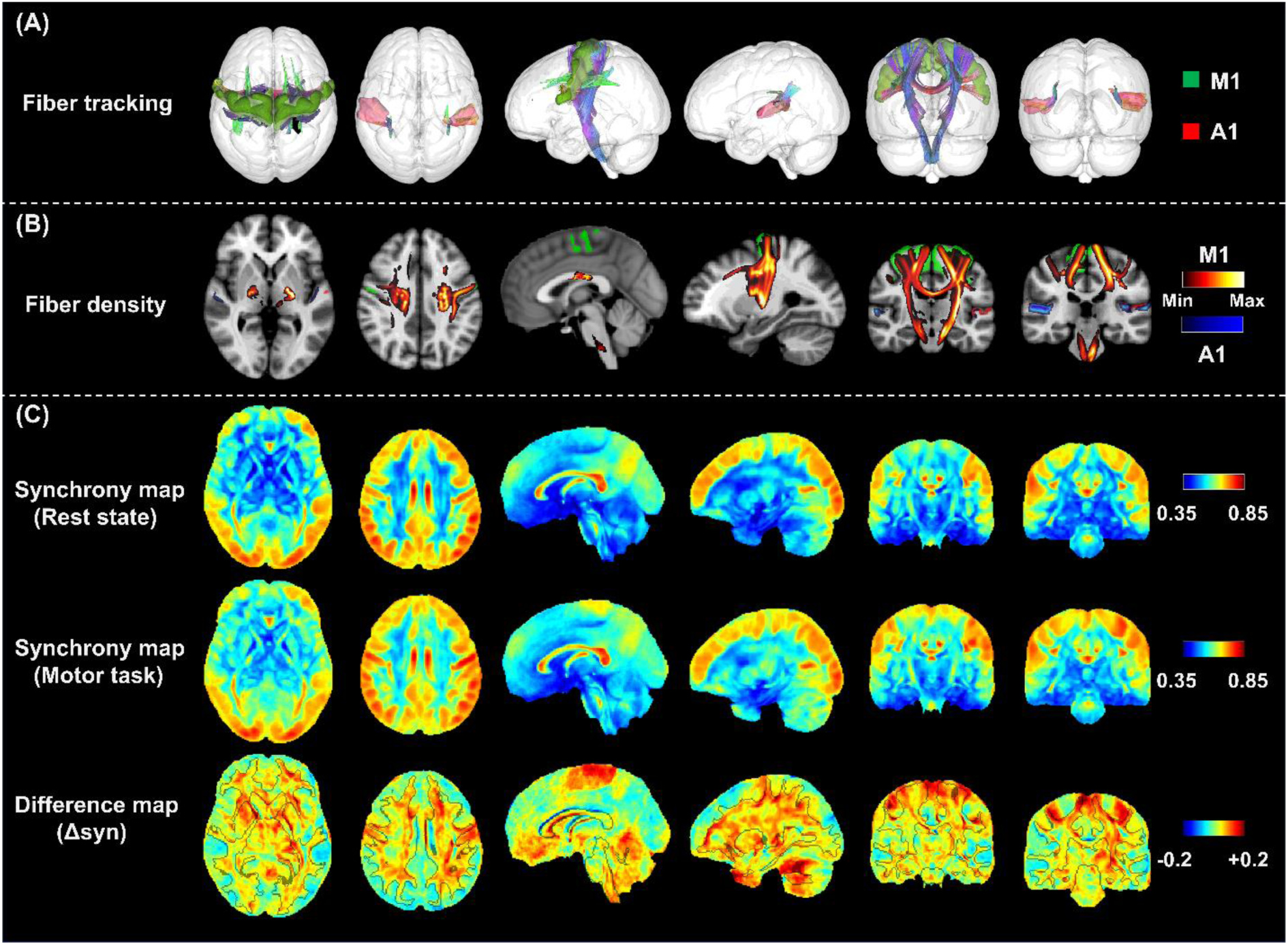
Comparison of group-average activation maps obtained from resting-state fMRI data and motor-task fMRI data on the basis of WM fiber tractograms. (A), exhibition of fiber tracking. The primary motor cortex (M1) and auditory cortex (A1&A2) related WM regions were determined by using fiber tracking with seed points defined in the corresponding cortical regions. (B), typical slices of fiber intensity maps estimated from fiber tracking. The fiber intensity of the M1 and A1&A2 related WM regions are displayed with red-yellow and blue colors, respectively. (C) synchrony maps and the corresponding difference maps displayed at the same slice indexes with the images in B. In both M1 GM regions and the related WM regions, the synchrony maps exhibit obviously higher intensities under motor task than those estimated from the resting-state, which demonstrates coactivations of the associated GM and WM regions. In addition, some WM regions connected with the occipital and frontal lobes also exhibit obvious enhancements of synchrony under the motor task whereas the WM regions associated with the auditory cortex have no significant changes between the two states.

**Fig. 4.**
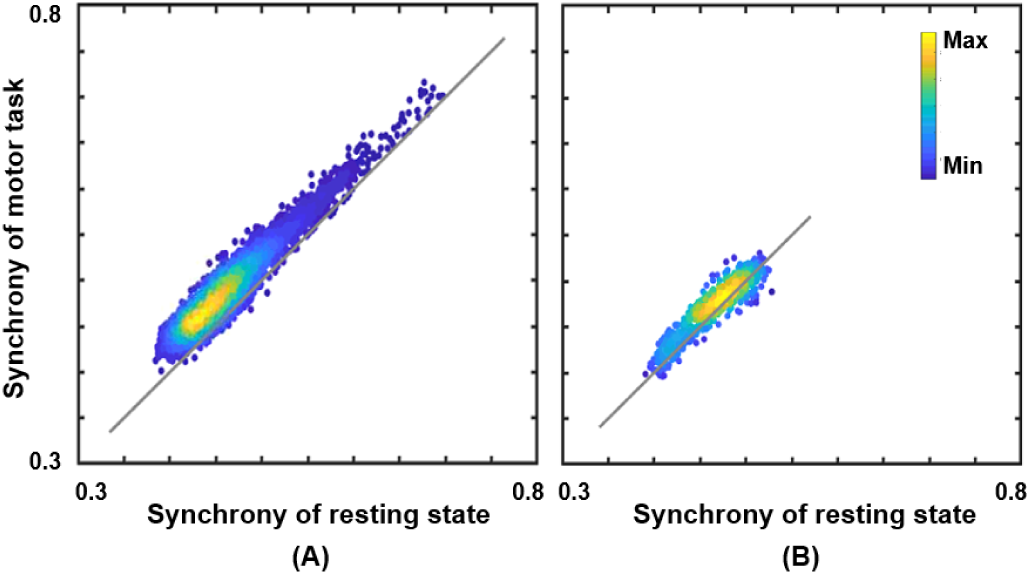
Scatter pots of activations for the voxels in the primary motor-cortex (M1) related WM region (A) and the auditory cortex (A1&A2) related WM region (B). Distances from points to an identity line measure the degree of activations. Activations in the M1 related WM region are significantly higher than those of the A1&A2 related WM region, in which the scatter points are tightly and symmetrically distributed along the identity line.

In **Fig. 5**, tract-wise changes in synchrony from resting state to motor task are examined for 46 fiber tracts defined in the HCP population-averaged tractography atlas. **Fig. 5A** shows statistical comparisons for each fiber tract, where paired student tests were used to identify significant changes in tract-averaged synchronies from 64 individual subjects. As seen, the synchrony during the motor task significantly increased for 33 fiber tracts with P < 0.05, and for 16 fiber tracts with P < 0.01. Among those tracts that changed, the majority are directly engaged in sensorimotor functions or visual processing including tracts in/linking the cerebellum and brainstem (cerebellum (CB), middle cerebellar peduncle (MCP), superior cerebellar peduncle (SCP), inferior cerebellar peduncle (ICP), vermis (V) and medial lemniscus (ML)) ^38,39^, the corticospinal tract (CST) ^40^, the corpus callosum (CC) ^41^ and the optic radiation (OR) ^42^. Note that many fiber tracts that are not directly involved in motor function or vision also show changes in **Fig. 5A** presumably because they are associated with motor network indirectly. For instance, the observed changes in the corticostriatal pathway (CS) may be explained by its role in motor planning as this fiber tract connects to the putamen, a main basal ganglia structure responsible for motor control ^43^. Tract-wise increases in synchrony are further visually illustrated in **Fig. 5B**. Here, the differences in tract-averaged synchrony are assigned as intensities to the corresponding tracts with significant increases in synchrony (P < 0.05), which are then displayed as 2D images using maximum intensity projection for axonal, sagittal and coronal views. High intensities in the figure appear mainly along the CST, consistent with the functional dominance of the sensorimotor pathways involved in the execution of the motor task.

**Fig. 5.**
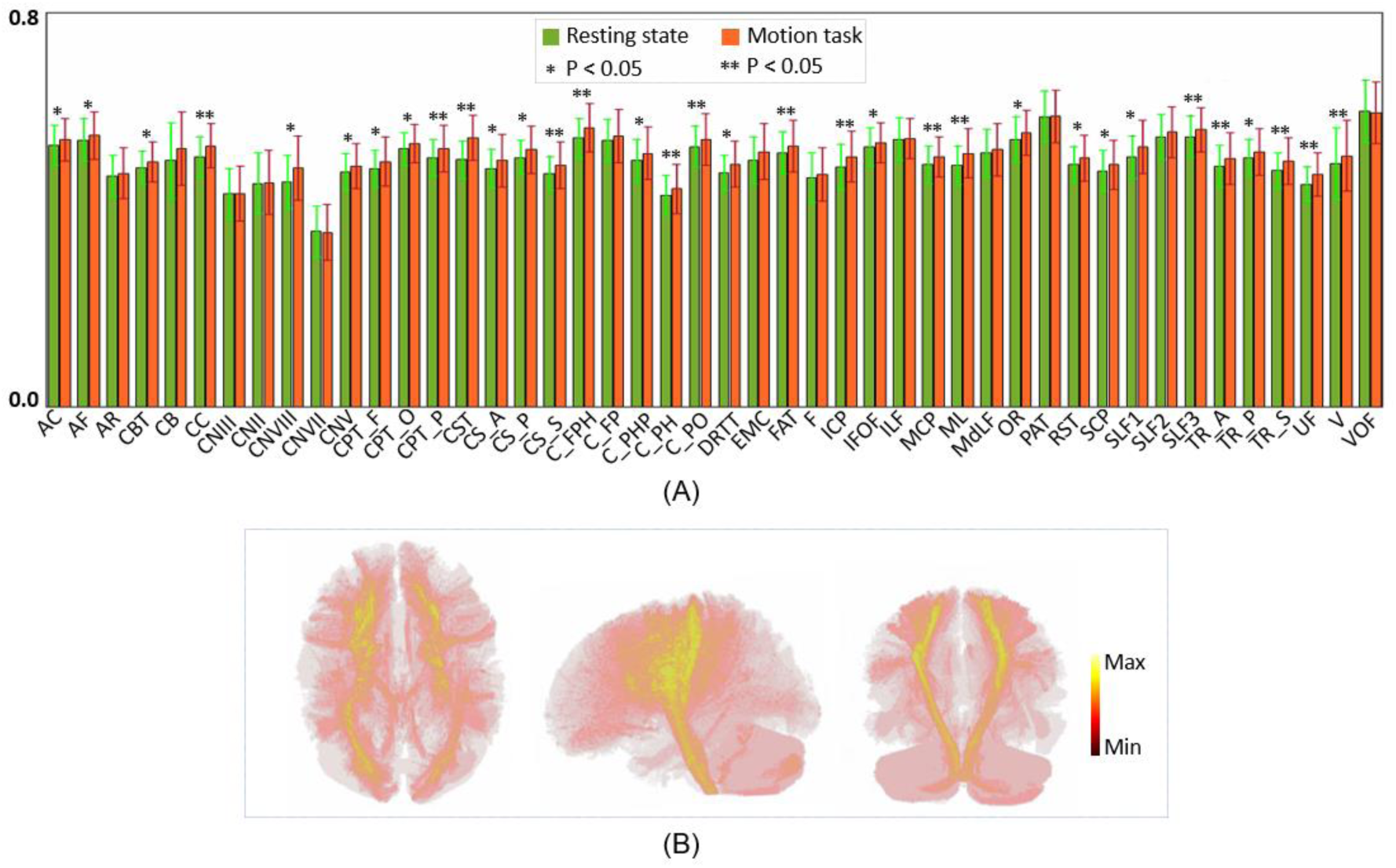
Tract-wise comparison of synchrony between resting state and motor task for 46 fiber tracts defined in the HCP population-averaged tractography atlas. (A), statistical comparisons of synchrony for each fiber tract. Paired t-test was implemented with tract-averaged synchronies estimated individually from 64 subjects. The result of statistical analysis shows that the synchrony under the motor task significantly increases for 33 fiber tracts with P < 0.05, and for 16 fiber tracts with P < 0.01. (B), maximum intensity projection images of differences in tract-averaged synchronies. To visually illustrate activations of the fiber tracts, the differences in tract-averaged synchrony are assigned as intensities to the corresponding tracts with significant increases in synchrony (P < 0.05), which are displayed as 2D images using maximum intensity projections for axonal, sagittal and coronal views. High intensities in the figure appear mainly along the CST, consistent with the functional dominance of the sensorimotor pathways involved in the execution of the motor task.

**Fig. 6** compares group level statistical maps of Δsyn-based and GLM derived changes (B), with FA color maps as anatomical references of WM (A). In the statistically inferred activation maps from both approaches, t-values thresholded with FDR-corrected P values of 0.05 and 0.01 are shown in **Fig. 6B**. The GLM-indicated activation maps show that the whole M1 cortical region is engaged by the composite task, that includes foot, hand and tongue movements, which agree well with previous results ^44^. Besides, the insular cortex that is engaged in motor control also exhibits activations in GLM statistical maps. The Δsyn-based results also highlight the cortical region of M1 with smaller coverages in the tongue motor cortex and insular cortex than the GLM-based activations. This can be actually deduced from **Fig. 3** in which these regions in Δsyn-based activation maps have similarly high synchronies in both the resting state and the motor task. On the other hand, there are significant Δsyn-based changes in WM regions that are hardly captured by the GLM approach. Theses detected activations span large WM regions that are directly or indirectly associated with M1 (See the preceding paragraph). This demonstrates the power of our approach to detect functional changes in WM, though some of the detected changes in WM appear to be more disperse or discontinuous than in GM. This phenomenon may largely be attributable to inter-subject variabilities of functional structures in WM, which typically have elongated geometry, thus tending to confound group level statistical analyses due to imperfect registrations. These spatial variabilities are demonstrated by the synchrony structures denoted by the red arrows in **Fig. 6C**, which shows synchrony maps of three typical subjects that have been registered into MNI space. In spite of the use of fine nonlinear registrations, elongated subtle synchrony structures in WM show discernable spatial variations among these subjects.

**Fig. 6.**
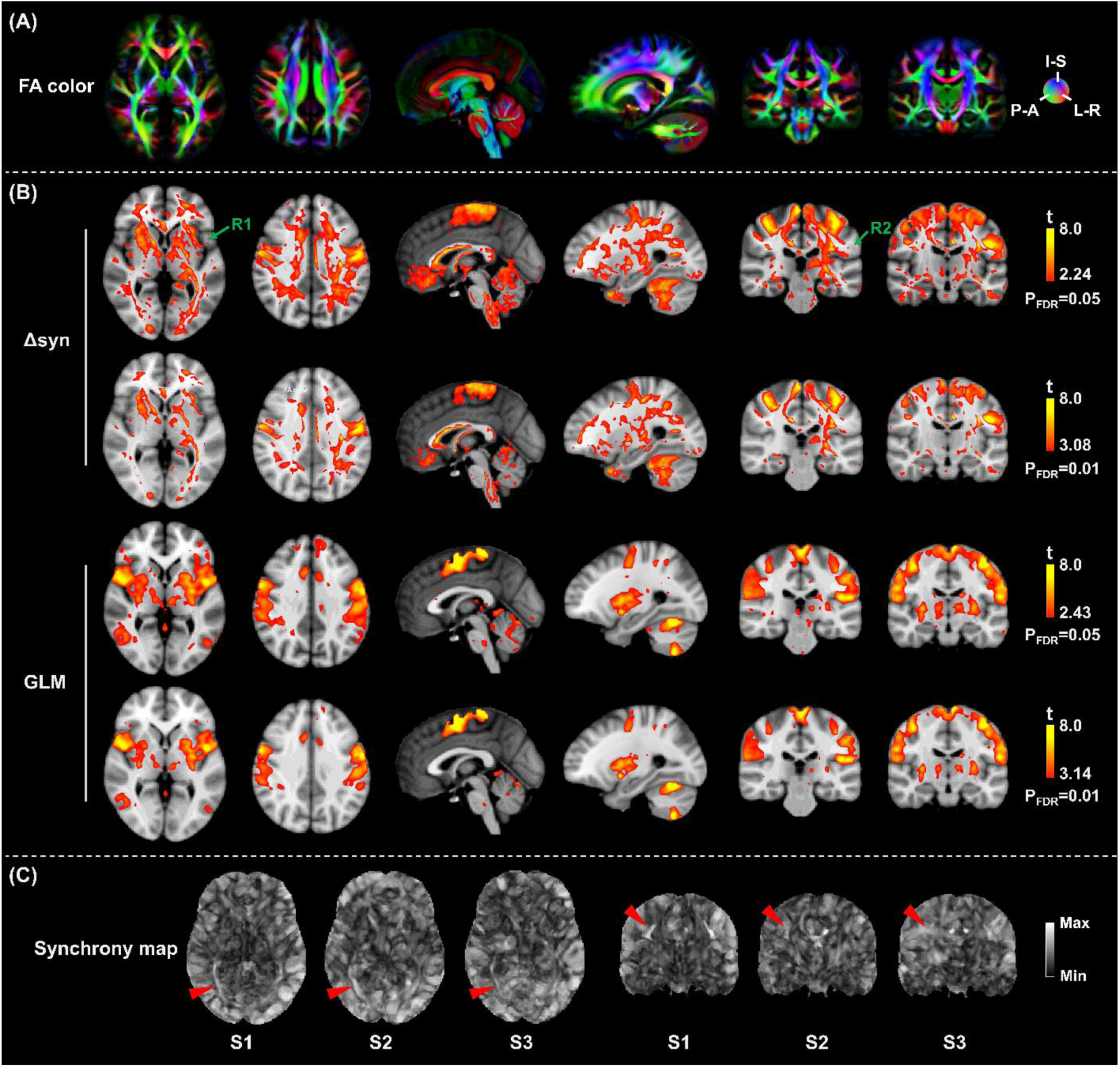
Comparison of group level statistical maps of Δsyn-and GLM-based activation with anatomical references of fiber structures. (A), FA color maps from one representative participant. The FA color maps show fiber structures. (B), group level statistical maps of Δsyn-and GLM-based activation. The statistical maps of Δsyn-based changes are derived by voxel-wise paired t-test of differences between resting state and motor task with a false discovery rate (FDR) correction for the 64-subject dataset. The statistical maps of GLM-based activation are estimated by using the standard SPM toolbox. In the statistically inferred activation maps from both the approaches, t-values thresholded with the FDR-corrected P values of 0.05 and 0.01 are shown in B. The GLM-based activation maps indicate that the whole primary motor cortex (M1) is engaged by the composite task that includes foot, hand and tongue movements. The Δsyn-based activations also highlight the cortical region of M1 and show smaller coverages in the insular cortex (R1) and the tongue motor cortex (R2) than the GLM-indicated activations. Moreover, the Δsyn-based activations exhibit in WM regions that are hardly captured by the GLM approach. The green arrows indicate the activations in WM that have similar spatial patterns with fiber structures identified in FA color maps. (C), synchrony maps of three typical subjects displayed in MNI space. There are discernable inter-subject variabilities of elongated fibers (indicated by red arrows) among these subjects, which could confound the statistical analysis at group level, and thus may account for the dispersity and discontinuity of activations detected in WM.

**Fig. 7** demonstrates the reproducibility of the FAIW-PCA activation mapping, which is validated by two independent datasets (G1 and G2). The group-average Δsyn maps estimated from the two datasets are displayed in **Fig. 7A**, which show very similar spatial patterns of the enhancement of synchrony by the task. **Figs. 7B&C** show statistical t maps of Δsyn-based activations for the two datasets, which are thresholded with FDR-corrected P values of 0.05 and 0.01. To quantify the similarity between the activation maps from the two datasets, the Dice coefficient was calculated as

**Fig. 7.**
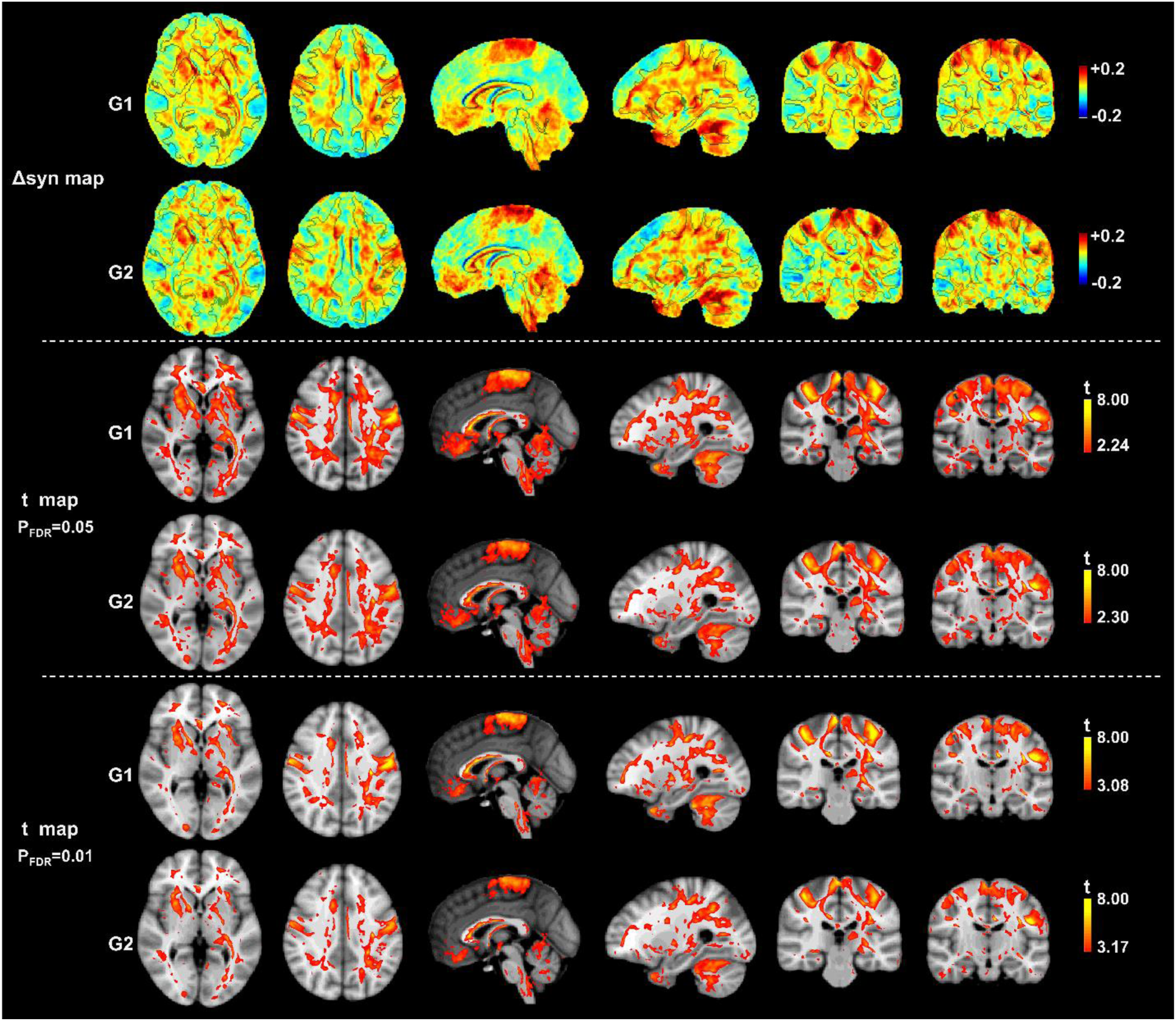
The reproducibility of the FAIW-PCA activation mapping validated with two independent datasets (G1 and G2). (A), group-average Δsyn maps estimated from the two datasets. The two maps exhibit similar spatial patterns of the task-induced enhancement of synchrony. (B&C), statistical t maps of Δsyn-based activation from the two datasets. In B&C, t values are thresholded with the FDR-corrected P values of 0.05 and 0.01, respectively. Dice coefficient is calculated to quantify the similarity between the activation maps from the two datasets. Dice coefficients of the activation maps estimated for B and C are 0.79 and 0.84, respectively, which demonstrates a good reproducibility of the proposed approach.

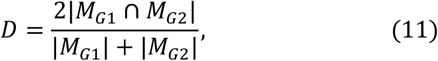

where *M*_*G*1_ and *M*_*G*2_ are two sets of activated voxels detected from G1 and G2, respectively and |·| denotes set cardinality. The Dice coefficient is constrained to the range of [0, 1], where a value 1 signifies perfect overlap between the two maps and a value 0 represents no overlap. In this study, Dice coefficients of the activation maps in B and C are 0.79 and 0.84, respectively, which demonstrates a high reproducibility of the FAIW-PCA activation mapping.

## IV. Discussion

Localizing neural activations under specific functional loadings is fundamental to understanding the functional organization of the brain. GLM-based activation mapping has been the standard approach for detecting and localizing task-evoked activations in GM, and assumes that BOLD signals in response to neural activation can be modeled by convolving the time course of neural activity with a fixed HRF that is determined a priori ^45^. However, this approach is inappropriate for WM. In this study, we proposed a model-free approach to detect changes in BOLD signals associated with neural activities in WM by measuring the enhancement of synchrony in WM BOLD signals during a functional task. The model-free approach draws upon an assumption that, under functional loadings, BOLD signals in relevant WM fibers are modulated by stimulus-evoked neural responses and thereby become more synchronized than in a resting state. This approach consists of two major steps. First, a fiber-architecture-informed spatial window is created with ODFs constructed from HARDI data, which characterizes a topographically defined neighborhood of a voxel that is isomorphic to local fiber architectures within WM. Second, a modified PCA is used to estimate the synchrony of BOLD signals based on the spatial window. The proposed approach is validated using a cohort of 3T fMRI datasets from the HCP, which demonstrates that the changes associated with neural activation can be reliably detected as enhancements of fMRI signal synchrony in the fibers that are engaged in the presented task.

Structures defined by intrinsic synchronous signals exist in both GM and WM and have similar boundaries with other definitions of anatomical and functional structures (**Fig. 2**). For GM, they appear as separable volumes with low degrees of anisotropy that are consistent with identifiable functional areas or units that to date have been implicitly assumed in the study of the functional connectome (i.e., Brodmann areas). In contrast to GM, functional units of WM measured by fMRI are elongated fibers that facilitate the transmission of neural signals along the fibers, which are identifiable by WM synchronous structures shown in **Fig. 2 and Fig. 5C** at group and individual levels, respectively. These synchronous components can be clearly observed in a resting state and become more pronounced in functionally-relevant structures under task loading (**Fig. 3**), which agrees with previous findings that correlations of BOLD signals in relevant WM bundles increase with tasks ^24-26^. Note that the regions that contain parallel fascicles tend to have higher levels of synchrony than those with incoherent orientations because some fibers in the latter scenario are in principle connected to different functional units in GM such that synchronous BOLD signals are not apparent.

By comparing group-average synchrony maps derived from resting-state and motor-task fMRI data, we demonstrate that BOLD signal synchrony can be enhanced by task loading in both GM and WM (**Fig. 3, 4 and 5**). For the GM, group level statistical maps of Δsyn-based activations in the M1 region show a moderate overlap with those of GLM-based activations, although with a smaller coverage than the latter, especially in the tongue motor cortex and the insular cortex (**Fig. 6**). The missing voxels of Δsyn-based activations in the tongue motor cortex may be explained by subconscious swallowing activity that inevitably occurs during the resting state acquisitions, which is implied by the observation in **Fig. 3** that the missing regions in Δsyn-based activation maps have similar synchronies in both the resting state and the motor task. Likewise, the insular cortex, which possesses multiple functions such as interoceptive attention, multimodal sensory processing, autonomic control and perceptual self-awareness ^46,47^, may be equally activate in the two states, and thus show similar synchronies. The Δsyn maps exhibit large increases in the M1-related WM regions in **Fig. 3**, which demonstrates that Δsyn-based activation mapping possesses a distinctive capability to detect activity-related signal changes in WM that are not captured by the GLM approach. This highlights the possibility that changes in synchrony do not depend on net increases in BOLD signal. In addition, some WM regions leading to the occipital and frontal lobes also exhibit obvious enhancements of synchrony under the motor task in **Fig. 3**. The spatial patterns of synchrony increases are consistent with previous reports that the occipital and frontal lobes are respectively engaged in visual processing ^48,49^ and executive functions ^50^ associated with the motor task. Further statistical analysis based on a population-averaged tractography atlas demonstrate that many fiber tracts that are indirectly associated with M1 also show significant enhancements of BOLD signal synchrony (**Fig. 5**). This implies a massive engagement of WM in the execution of the motor task, which agrees with earlier reports that GM across almost the entire brain can be activated by a simple task. Although many WM regions are involved in the motor task ^51^, the M1-related WM exhibit most significant changes, as shown **Fig. 5B**.

In this study, the sensitivity, specificity and reproducibility of the proposed approach have been comprehensively investigated. First, the Δsyn-based activations can be clearly observed in the M1-related WM regions, suggesting this approach has reasonable sensitivity. It is well-known that the magnitudes and signal-to-noise ratios of BOLD signals in WM are substantially lower than those in GM [19] due to the significantly lower blood volume in WM compared to that in GM [1, 20], so detecting changes in the magnitudes of BOLD signals in a task is challenging. Alternatively, our model-free approach exploits the enhancements of BOLD signal synchrony to identify neural activations, which is robust in the presence of noise without modeling the BOLD signal. Second, the specificity of the proposed approach is only partially validated by the observation that there are no significant enhancements of synchrony in auditory cortex under the motor task (**Fig. 4B**). We found that some other WM regions that are not expected to be directly associated with the task targeted M1 region also show significant enhancements of synchrony, though these may be indirectly affected. Third, the reproducibility of the proposed approach was validated using two independent groups of participants where statistical t maps of Δsyn-based activation thresholded with the FDR-corrected P values of 0.05 and 0.01 were compared. Dice coefficients of the activation maps estimated from the two groups are 0.79 and 0.84, respectively, which demonstrates high reproducibility of the FAIW-PCA activation mapping.

To validate the proposed approach with large-sample datasets, we analyzed the open-access HCP datasets of 128 participants, though the parameters of MRI acquisition and the designs of the fMRI tasks were not tailored for our study, which thus led to some limitations. First, parameters of MRI acquisitions in the HCP data are optimized for fMRI of GM. In a future study, the parameters of MRI acquisitions can be optimized for detecting synchrony of BOLD signals in WM, which may not require high temporal resolution. Second, the motor task of the HCP involves five different motions that produce only about 18 data points in each time courses for each motion. Some regions in brain that are engaged by only one motion may be more difficult to be detected by our approach because the measurement of the synchrony is sensitive to the number of data points in time courses. This limitation in the task design can reduce the sensitivity of the proposed approach, which may be one reason for the loss of activations detected compared to the GLM results. In a future study, increasing the data points that are modulated by identical repeated stimuli can be used to increase the sensitivity of the proposed method.

## V. Conclusions

The GLM-based activation mapping that is well-established for detecting BOLD signal changes in a task in GM is not well suited for studies of activity-related signal changes in WM. In this study, we proposed a model-free approach to detect task-evoked signal changes in WM by measuring increases in BOLD signal synchrony within local fiber architectures. Using the HCP datasets, we validated that the proposed approach could detect BOLD signal changes in WM fibers that engaged in the motor task with high sensitivities and reproducibility. This approach promises to provide a standardized tool to map neural responses in WM in the future study.

## Declaration of competing interests

The authors declare that they have no competing interests.

## Code and data availability

An implementation of the approach proposed in this work will be made available as a MATLAB package on GitHub. Customized links will be added in the final version of the manuscript. Human MRI datasets used in this study were sourced from the WU-Minn Human Connectome Project (HCP) database.

## Acknowledgements

Data used in this work were provided by the Human Connectome Project, WU-Minn Consortium (Principal Investigators: David Van Essen and Kamil Ugurbil; 1U54MH091657) funded by the 16 Institutes and Centers of the National Institutes of Health (NIH) that support the NIH Blueprint for Neuroscience Research; and by the McDonnell Center for Systems Neuroscience at Washington University. This work was supported by National Institutes of Health under Grant R01NS093669 and Grant R01NS113832.

## Notes

### Competing Interest Statement

The authors have declared no competing interest.

### Summary of Updates

An author has been added.

